# Cell cycle variants during *Drosophila* accessory gland development

**DOI:** 10.1101/719013

**Authors:** Allison M. Box, Navyashree A. Ramesh, Shyama Nandakumar, Samuel Jaimian Church, Dilan Prasad, Ariana Afrakhteh, Russell S. Taichman, Laura Buttitta

**Affiliations:** Department of Molecular, Cellular and Developmental Biology, University of Michigan, Ann Arbor, MI 48109, USA; Department of Periodontology, School of Dentistry, University of Alabama at Birmingham, Birmingham, AL; University of Pittsburgh, Department of Cell Biology, Pittsburgh, PA, USA

## Abstract

The *Drosophila melanogaster* accessory gland is a functional analog of the mammalian prostate containing two secretory epithelial cell types, termed main and secondary cells. This tissue is responsible for making and secreting seminal fluid proteins and other molecules that contribute to successful reproduction. The cells of this tissue are bi-nucleate and polyploid, due to variant cell cycles that include endomitosis and endocycling during metamorphosis. Here we provide evidence of additional cell cycle variants in this tissue. We show that main cells of the gland are connected by ring canals that form after the penultimate mitosis and we describe an additional post-eclosion endocycle required for gland maturation that is dependent on juvenile hormone signaling. We present evidence that the main cells of the *Drosophila melanogaster* accessory gland undergo a unique cell cycle reprogramming throughout organ development that results in step-wise cell cycle truncations culminating in cells containing two octoploid nuclei with under-replicated heterochromatin in the mature gland. We propose this tissue as a model to study developmental and hormonal temporal control of cell cycle variants in terminally differentiating tissues.

## Introduction

The *Drosophila* accessory gland is functionally analogous to the mammalian prostate. This tissue is an essential component of the male reproductive system and is responsible for making and secreting seminal fluid proteins, sex peptides, and anti-microbial proteins that are transferred to the female upon mating (Adams & Wolfner, 2007; Heifetz et al., 2005; Lung et al., 2001; Qazi & Wolfner, 2003; Ravi Ram et al., 2005) The accessory gland consists of two lobes, each is comprised of a single layer of secretory epithelial cells that form a large lumen apically, and are surrounded basally by extracellular matrix and enclosed in a muscle layer (Bairati, 1968; Susic-Jung et al., 2012). Each lobe of the accessory gland consists of approximately 1,000 epithelial cells, which are made up of two cell types: main cells and secondary cells (Bairati, 1968). Main cells are the smaller of the two types and are hexagonal in shape. These cells make up a majority of the gland and are located mostly in the proximal and medial portions of the lobes. Secondary cells are larger, more luminal cells that are located at the distal tip of the lobes. There are approximately 40-60 secondary cells in each lobe, with the rest of the cells (∼940-960 cells) being main cells. Main and secondary cells have distinct but partially overlapping secretory protein profiles and both cell types play an important role in fecundity (Bertram et al., 1992; Sitnik et al., 2016).

Previous work shows that accessory gland development takes place during larval and pupal stages and undergoes tight cell cycle regulation that includes variant cell cycles. In late larval stages, FGF signaling drives recruitment of mesodermal cells to the genital disc. These cells undergo a mesenchymal to epithelial transition and give rise to the precursors for the accessory glands and seminal vesicles (Ahmad & Baker, 2002). During early metamorphosis, the accessory gland progenitors increase in number by standard mitotic cell cycles. Around 50-55 hours after pupa formation, the cells of the developing accessory gland arrest proliferation and synchronously enter a truncated, variant cell cycle, in which nuclear division occurs but cytokinesis does not, resulting in the bi-nucleation of the epithelial cells (Taniguchi et al., 2014). Approximately 10 hours later, the cells enter an additional synchronized endocycle, increasing their DNA content (C) without mitosis (Taniguchi et al., 2014). This results in ∼1,000 8C binucleated cells, each cell containing two 4C nuclei (Taniguchi et al., 2014). It has been previously thought that after the endocycle in the pupal stage the cells of the accessory gland exit the cell cycle, however more recent studies have shown that secondary cells retain the capacity to increase DNA content in response hormonal or mating signals (Leiblich et al., 2019).

In this study, we closely examine the cell cycle status of the main cells in the adult accessory gland. Starting immediately post-eclosion we find evidence of a previously undescribed, tissue-wide endocycle in the newly eclosed adult main cells. Induction or reduction of cell cycle and growth regulators is sufficient to alter the level of endocycling in the adult gland. Additionally, like many other endocycles, we observe that heterochromatin is underreplicated and delayed in a manner controlled by G1 Cyclin/Cdk activity in the adult main cell endocycle. We show that juvenile hormone signaling is required for this adult-stage endocycle and disruption of this signaling pathway can affect proper tissue development and fertility. Additionally, we see evidence of ring canals in the adult tissue, suggesting that an additional variant cell cycle occurs prior to what has been previously described during the pupal stage (Taniguchi et al., 2014). Altogether, our findings establish that the *Drosophila* accessory gland main cells undergo developmental reprogramming of the cell cycle via tightly regulated, step-wise cell cycle truncations, resulting in a uniquely polyploid and bi-nucleate secretory organ.

## Results

### Early growth and endocycling of main cells in the adult *Drosophila* accessory gland is conserved

As a result of variant, truncated cell cycles that occur during pupal development, the epithelial cells of the adult *Drosophila* accessory gland (AG) are polyploid and binucleated (Taniguchi et al., 2014) (Figure 1A). Like the mammalian prostate, the fly AG grows with age and AGs from virgin males undergo a period of rapid growth from the day of eclosion to day 10 post-eclosion (Figure 1B, E).

**Figure 1:**
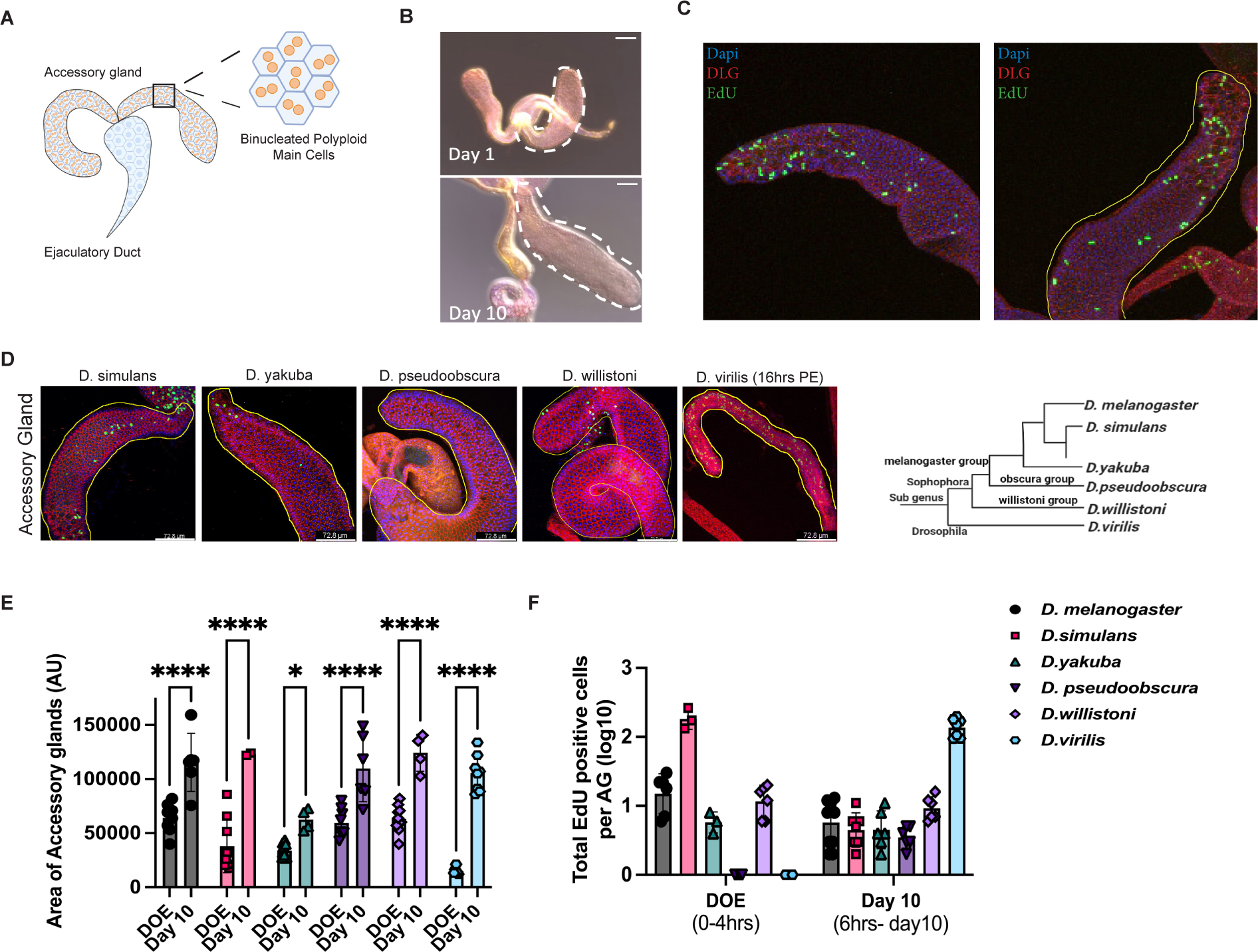
Early growth and the endocycling of main cells in the adult *Drosophila* accessory gland is conserved. A: Cartoon representation of the male reproductive gland. Boxed area represents area in which nuclear ploidy measurements were obtained in the adult accessory gland for this manuscript, zoom shows the structure of main cells, including the binucleation. B. Images of adult accessory glands on the day of eclosion and day 10 post eclosion. Accessory gland lobe is outlined with white dashed line. The adult tissue undergoes dramatic growth between eclosion and day 10. C: EdU incorporation in newly eclosed male accessory glands. Dissection occurred between 1-3 hours post eclosion and labeling was *ex vivo* for 1 hour in Ringers. Accessory gland lobe is outlined with yellow line. D: EdU incorporation in newly eclosed male accessory glands in other species of *Drosophila*. Dissection occurred between 1-3 hours post eclosion, with the exception of *D. virilis* which was dissected at 16 hours post eclosion. Labeling was ex vivo for 1 hour in ringers. Accessory gland lobe is outlined with yellow dashed line. E: Quantifications of gland area for *D. melanogaster* and other species of *Drosophila* on the day of eclosion (DOE) and Day 10 (virgin). Gland growth between DOE and Day 10 is conserved. F: Quantifications of EdU positive cells per AG (log10) for many species of *Drosophila* on the day of eclosion (DOE) and Day 10 (virgin). The left grouping shows number of cells positively labeled during a 1 hour labeling on DOE and the right grouping shows the number of cells positively labeled throughout a 10 day feeding experiment where animals are fed EdU+Sucrose from 6 hours post eclosion to dissection. EdU incorporation is conserved across all species analyzed. Statistical Analysis Performed:

While a majority of AG growth in early adulthood can be attributed to the production and secretion of sex peptides, which expands the lumen of the gland, we hypothesized that changes in cell number or cell size may also support this period of rapid growth. To address whether changes in cell number contribute to gland growth, we stained the adult AG for the mitotic marker, phospho-histone H3 (PH3), at various time points throughout the male lifespan. In sum, over 100 AGs, of various ages and mating status, were negative for PH3 staining, while cell size and nuclear size increase with age (Sup Figure 1A), suggesting that under normal physiological conditions, gland growth is coupled to changes in cell size rather than cell number.

Many *Drosophila* tissues undergo a variant cell cycle called an endocycle in order to increase tissue size in the absence of mitosis under normal physiological conditions (Edgar et al., 2014; Orr-Weaver, 2015). Endocycling occurs when cells modify the canonical cell cycle to cycle through G/S phase without entering an M phase, thereby increasing DNA content (C) of a cell, resulting in polyploidy. Increased ploidy can contribute to increased biosynthesis, as is the case with *Drosophila* nurse cells (Inge Øvrebø & Edgar, 2018; Orr-Weaver, 2015) and recently the secondary cells of the adult AG have been shown to endocycle in response to mating (Leiblich et al., 2019). As the adult AG is highly secretory and responsible for making large amounts of AG specific proteins and other molecules for transfer to the female upon mating (Prince et al., 2019; C. Wilson et al., 2017), we examined whether DNA replication also occurs in the main cells of the adult AG.

To visualize cells that have undergone S-phase or DNA synthesis, we first fed adult flies with 5-Ethynyl-2′-deoxyuridine (EdU) for 10 days post-eclosion and observed a low level of EdU incorporation in main cells demonstrating that on average about 3-4 main cells per gland endocycle at least once during the first 10 days of adulthood (Sup Figure 1B). In these EdU ingestion assays, we used blue dye to ensure we only analyzed the animals that had consumed the EdU mixture. During these assays, we noted that male flies do not eat for the first 5-8 hours post-eclosion (Sup Figure 1C, D). We therefore decided to examine the AG at timepoints immediately post-eclosion, before the EdU molecule is ingested. We dissected and directly exposed tissues in Ringer’s solution with EdU for one hour. We observed that 100% of glands exhibit widespread EdU labeling in a large fraction of the main cells, suggesting that the adult AG undergoes an early, relatively synchronous round of DNA replication immediately post-eclosion (Figure 1C, Figure 2A). We believe this additional endocycle in the early adult stage has been previously missed due to the use of feeding-based EdU labeling assays in adult animals.

**Figure 2:**
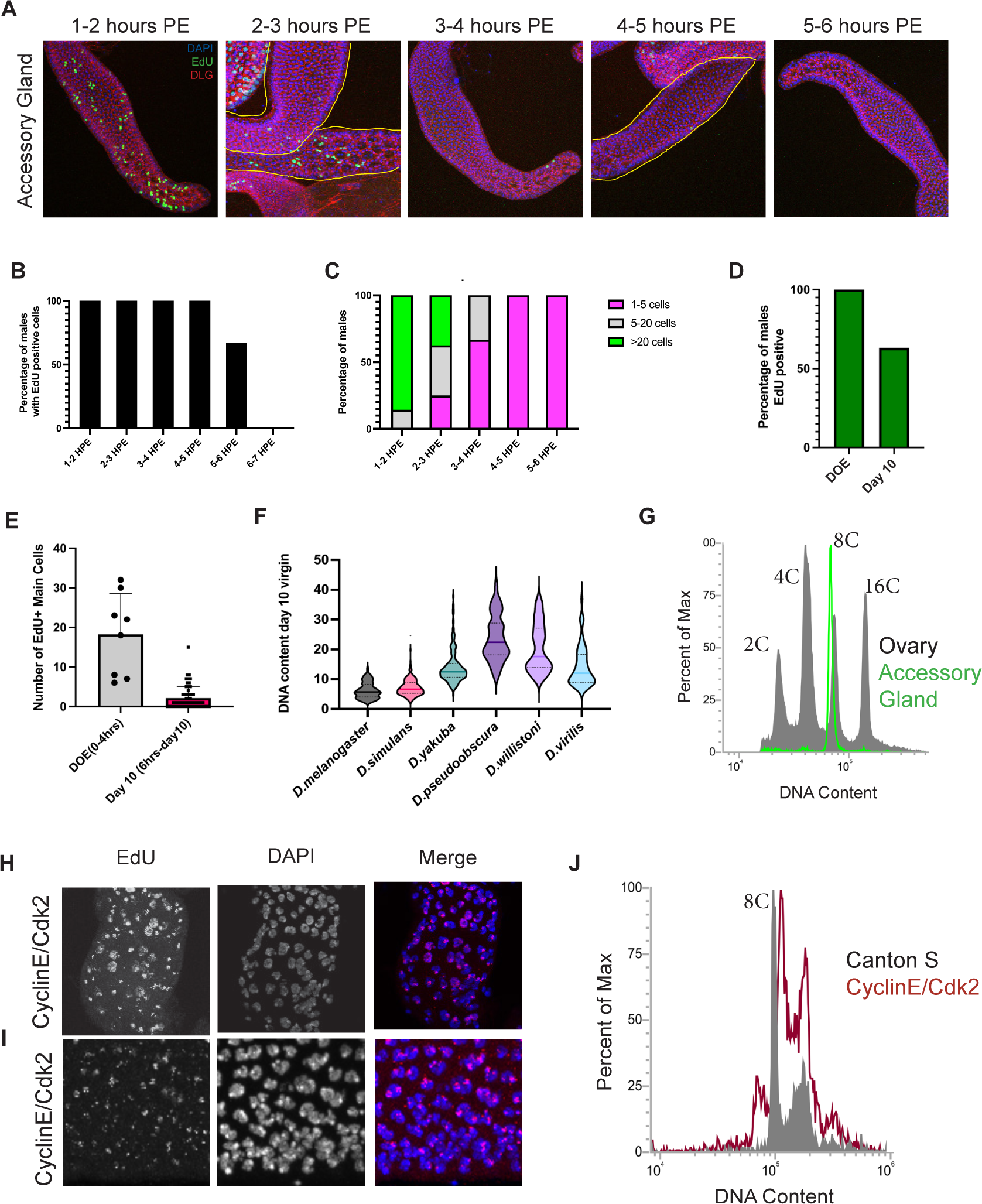
*Drosophila melanogaster* accessory gland main cells exit the cell cycle shortly post eclosion with 2 under-replicated 8C nuclei. A: EdU incorporation on the DOE as a time course. Ex vivo EdU labeling was done in the adult accessory gland from 1-6 hours post-eclosion in 1 hour increments. Most EdU incorporation occurs within the first 3 hours post eclosion. B: Quantification of the percentage of males showing EdU positive cells in the accessory gland for each hour post eclosion during the time course above. EdU incorporation stops by 6 hours post eclosion. C: Quantification of the number of cells labeled in each accessory gland for each hour post eclosion during the time course shown above. The number of cells labeled is divided into 3 groups: 1-5 cells, 6-20 cells, and >20 cells. Levels of EdU incorporation drops dramatically by 3 hours post-eclosion. D: Quantification of the percentage of animals that are EdU positive on the day of eclosion (DOE) and Day 10. DOE incorporation is a 1 hour ex vivo labeling and Day 10 is a 10 day feeding of Sucrose+EdU. The feeding assay does not label cycling cells during first 6 hours post-eclosion due to time spent without eating immediately after eclosion (supplement). E: Quantifications of the number of EdU positive main cells on the day of eclosion (DOE) and Day 10. DOE incorporation is a 1 hour ex vivo labeling and Day 10 is a 10 day feeding of Sucrose+EdU. F: Ploidy measurements from mid-lobe main cells of Day 10 virgin males for multiple species of *Drosophila*. Ploidy of accessory gland main cells varies from specie to specie with *D. simulans* being the most similar to *D. melanogaster*. G: Flow cytometry histogram of nuclear DNA content from Day 5 virgin males. Grey plot shows polyploid control using Canton S ovaries and green plot shows nuclear DNA content from adult accessory glands. The accessory gland plot is shifted slightly to the left of the plot from the ovary, suggesting that the DNA is under-replicated. H and I: EdU labeling on the day of eclosion accessory glands with CyclinE/Cdk2 overexpression using Prd-gal4. EdU is a 1 hour ex vivo labeling. Punctate EdU signal aligns with DAPI bright spots or heterochromatic regions, suggesting that the ectopic expression of CyclinE/Cdk2 can push replication of under-replicated regions. J: Flow cytometry histogram of DNA content. Grey plot shows RFP+ accessory gland control nuclei and red plot shows RFP+ accessory cells where CyclinE/Cdk2 is being overexpressed. The shift to the right suggests that CyclinE/Cdk2 overexpression can push under-replicated regions of the DNA to finish replication.

We next wondered whether this endocycle in the adult AG was strain or species specific. We analyzed AGs across several *Drosophila* species for rapid, early tissue growth (as well as other commonly used control *D. melanogaster* strains such as *w^1118^*) and performed EdU incorporation assays to assess if early adult main cell endocycling is a conserved phenomenon. We see significant tissue growth and evidence of S-phase between day of eclosion and Day 10 via EdU in all species we tested (Figure 1D, E, F). It is important to note though that the development, sexual maturation, and lifespan of these species are on a different timescale from *Drosophila melanogaster* (Ma et al., 2018; Markow & O’Grady, 2008), which may alter the timing of the early endocycle. This is most clear in *Drosophila virilis*, where EdU incorporation is not seen during the first four hours post-eclosion labeling, but instead widespread EdU incorporation in main cells occurs at 16 hours post-eclosion (Figure 1D, F).

### *Drosophila melanogaster* accessory gland main cells exit the cell cycle shortly post-eclosion with two under-replicated 8C nuclei

Due to the high levels of EdU incorporation we see within a one-hour labeling period, we next asked when the early adult endocycle is completed. We collected virgin males once an hour and performed 1-hour EdU labeling intervals up to 5 hours post-eclosion (i.e. labeling takes place 1-2 hours, 2-3 hours, 3-4 hours post-eclosion, etc). We see that a majority of EdU incorporation occurs within the first 3 hours post-eclosion for *D. melanogaster* (Figure 2A, B, C). While the percentage of animals positive for EdU incorporation doesn’t drop until 5 hours post eclosion, the number of EdU positive cells is greatly reduced by 3h post-eclosion. (Figure 2C). This suggests that in *D. melanogaster* the post-eclosion endocycle is highly synchronized, similar to the endocycle during pupal development.

To test whether there may be additional endocycles after the first wave within 3-4h post-eclosion, we fed virgin males an EdU+Sucrose mixture for various intervals up to 10 days post-eclosion, since this will not capture the post-eclosion endocycle which occurs before the males eat at 6h but will capture any later endocycles. Interestingly, we do see some EdU incorporation in most animals indicating most adult AGs do endocycle, but the number of cells per gland is very low, ranging from 5-12 over 10 days (Figure 2D, E), suggesting that only a small subset (∼1% of main cells) of cells may either delay their post eclosion endocycle or undergo an additional endocycle after the first post-eclosion wave resulting in a small number of 16C nuclei (32C cells). This demonstrates that most of the adult main cell endocycling occurs within the first few hours of eclosion prior to feeding. Prior studies of AG development used DNA quantification by propidium iodide or H2Av-RFP in the pupal and adult glands and reported that main cell nuclei reach 4C DNA content (2nucleiX4C=8C cells) shortly before eclosion(Taniguchi et al., 2012, 2018). The widespread post-eclosion EdU incorporation we observe suggests that a subset of, if not all, main cell nuclei in the adult tissue will become 8C shortly post-eclosion, making the ploidy of main cells in the adult tissue 16C (2X8C=16C). We therefore used multiple methods to re-examine the nuclear DNA content of the adult main cells after the wave of post-eclosion endoreplication to determine their ploidy.

We first performed DAPI intensity measurements on the main cell nuclei of the adult AG at 10 days post-eclosion. We used basally located muscle nuclei of the adjacent tissue, the ejaculatory duct, as our diploid DNA content control. Integrated DAPI intensity measurements of similar sized z-stacks of 10 day old AG main cell nuclei confirm that most main cell nuclei are ∼8C resulting in most main cells having a total ploidy of 16C (Figure 2F). We next performed 10 day EdU+Sucrose feeding and measured the DNA content of AG main cell nuclei across other *Drosophila* species and we find that main cell ploidy is quite variable across species, with *D. simulans* most closely resembling the ploidy observed in *D. melanogaster* (Figure 2F, Sup Figure 2A-H).

We further verified our ploidy measurements in *Drosophila melanogaster* AGs by performing flow cytometry DNA content analysis on isolated nuclei. Using the *Drosophila melanogaster* ovary as a reference for DNA content, we see that the AG nuclei of 10 day old virgin males exhibit ploidies near the 8C peak of the ovary. We also see a small population of AG nuclei near the 16C peak (Figure 2G), consistent with our 10day EdU feeding result (Figure 2D,E, Sup Figure 1B). This suggests that nearly all nuclei in the *D. melanogaster* AG undergo a post-eclosion endocycle transitioning from 4C to approximately 8C and that small subset of cells undergo an additional endocycle by day 10 post-eclosion becoming approximately 16C.

We noted that the 8C and 16C DNA peaks for adult AGs consistently appear to be shifted slightly to the left of the 8C and 16C peaks for the ovary (Figure 2G). This suggested to us there may be under-replication occurring during the early post-eclosion endocycle. This is reminiscent of the polyploid follicle cells of the *Drosophila* ovary, which have been described to have early and late replicating regions, as well as Drosophila salivary gland nuclei which under-replicate heterochromatin (Hua & Orr-Weaver, 2017; Nordman & Orr-Weaver, 2012; Spradling & Orr-Weaver, 2003; Yarosh & Spradling, 2014).

Typically, heterochromatic regions are replicated late in S-phase in mitotic cells (Leach et al., 2000). While they are often underreplicated in polyploid cells, prolonged expression of Cyclin E during S-phase or ectopic induction of Cyclin E can push these late heterochromatic regions to replicate in polyploid nuclei (Calvi et al., 1998; Lilly & Spradling, 1996).We therefore examined whether ectopic expression of CycE/Cdk2 would induce heterochromatic endoreplication in the main cells of the adult AG. We used *Prd-Gal4*, a driver that is specific to the AG in the male reproductive system (Sharma et al., 2017), to manipulate gene expression in this tissue. Overexpression of CycE/Cdk2 in the AG resulted in a distinct pattern of EdU incorporation appearing as bright foci in the heterochromatic region of most nuclei (Figure 2H, I) similar to what is seen when Cyclin E is prolonged or ectopically expressed in the ovarian nurse cells (Calvi et al., 1998; Lilly & Sptadling, 1996). Further, when we overexpress CyclinE/Cdk2, the 8C and 16C peaks of the *Drosophila* AG shift slightly to the right when compared to the AGs wildtype males, suggesting that this genetic manipulation induces replication of heterochromatin in this tissue (Figure 2J).

Altogether these data reveal an adult-stage, synchronous endocycle with a truncated S-phase resulting in under-replicated heterochromatin within 3 hours post-eclosion. We hypothesize that this widespread endocycle acts as an additional developmental stage for the AG that likely supports the extensive growth reported during the first few days of adulthood and tissue maturation.

### The E2F/Rbf Network and APC/C oscillations regulate adult stage main cell endocycles

We next investigated approaches to manipulate endocycling in main cells. Previous work has established an important role for the E2F/Rb network in *Drosophila* endocycling cells (Bosco et al., 2001; Zielke et al., 2011). In brief, the *Drosophila* E2F network consists of two E2Fs: E2f1, which is described to act as a transcriptional activator, and E2F2, which is described to act as a transcriptional repressor. DP is the common heterodimeric binding partner of both E2Fs. Most cycling and endocycling cells exhibit oscillatory E2F1 activity that increases transcription of genes that promote S-phase entry, including Cyclins/Cyclin-dependent kinases (Cdks) and factors that drive DNA synthesis, followed by S-phase dependent degradation (Zielke et al., 2011). Rbf is the *Drosophila* version of Retinoblastoma, which inhibits E2F activity and acts as a brake for E2F oscillations. This brake is released by phosphorylation and inactivation of Rbf by Cyclin D or Cyclin E paired with Cdks.

To characterize the novel endocycle that occurs shortly post-eclosion, we used *Prd-Gal4* to either knockdown or overexpress positive and negative regulators of the cell cycle in order to assess the impact of endocycle manipulations on main cell ploidy. To assess effects on endoreplication we used two experimental procedures, (1) a 1 hour labeling where dissected AGs from males 0-3 hours post-eclosion were incubated in Ringers with EdU to visualize changes in EdU incorporation compared to control and (2) DAPI fluorescence intensity measurements to analyze DNA content at day 10 post-eclosion. The combination of these experiments allows us to determine if a loss or increase of EdU on the day of eclosion represents a true loss or increase of endocycling rather than a delay in cell cycle entry or a slowing down of the endocycle.

We first depleted Rbf in the AG using RNAi and observed an increase of EdU incorporation on the day of eclosion as well increased DNA content at Day 10, with nearly half of nuclei measuring 16C, making the overall DNA content of main cells 32C (Figure 3A, C). This suggests Rbf acts to limit post-eclosion endocycling. To confirm this, we overexpressed Rbf280, a phospho-mutant version of Rbf that can no longer be inhibited by CycD or CycE -dependent kinase activity and therefore would be expected to block adult endocycles (Xin et al., 2002). We find that overexpression of Rbf280 resulted in two distinct phenotypes in main cells. In most cases, we saw decreased EdU incorporation shortly post-eclosion and nuclei with 4C DNA content on day 10, resulting in a majority of main cells with only 8C DNA content, suggesting that Rbf280 blocked the widespread endocycle as expected (Figure 3A, D). However, Rbf280 overexpression occasionally resulted in a second, unexpected phenotype with increased EdU labeling and higher nuclear DNA content at Day 10 where a majority of nuclei contain 16C DNA content, resulting in 32C main cells (Sup Figure 3A,B). Our data suggests that Rbf likely limits endocycling in the adult AG, however, as we have observed in wings expressing Rbf280 (Flegel et al., 2016), there can be compensatory mechanisms that increase cell cycling when Rbf suppression of E2F activity hits a certain threshold. To further confirm that Rbf acts to limit endocycling in adult glands, we overexpressed Cyclin D with its partner CDK4, which phosphorylates and inhibits endogenous Rbf in the AG. We observed a widespread increase in S-phases on the day of eclosion (Figure 3A) and increased ploidies at day 10 with nuclei up to 32C, resulting in bi-nucleate cells with DNA content up to 64C (Figure 3C).

**Figure 3:**
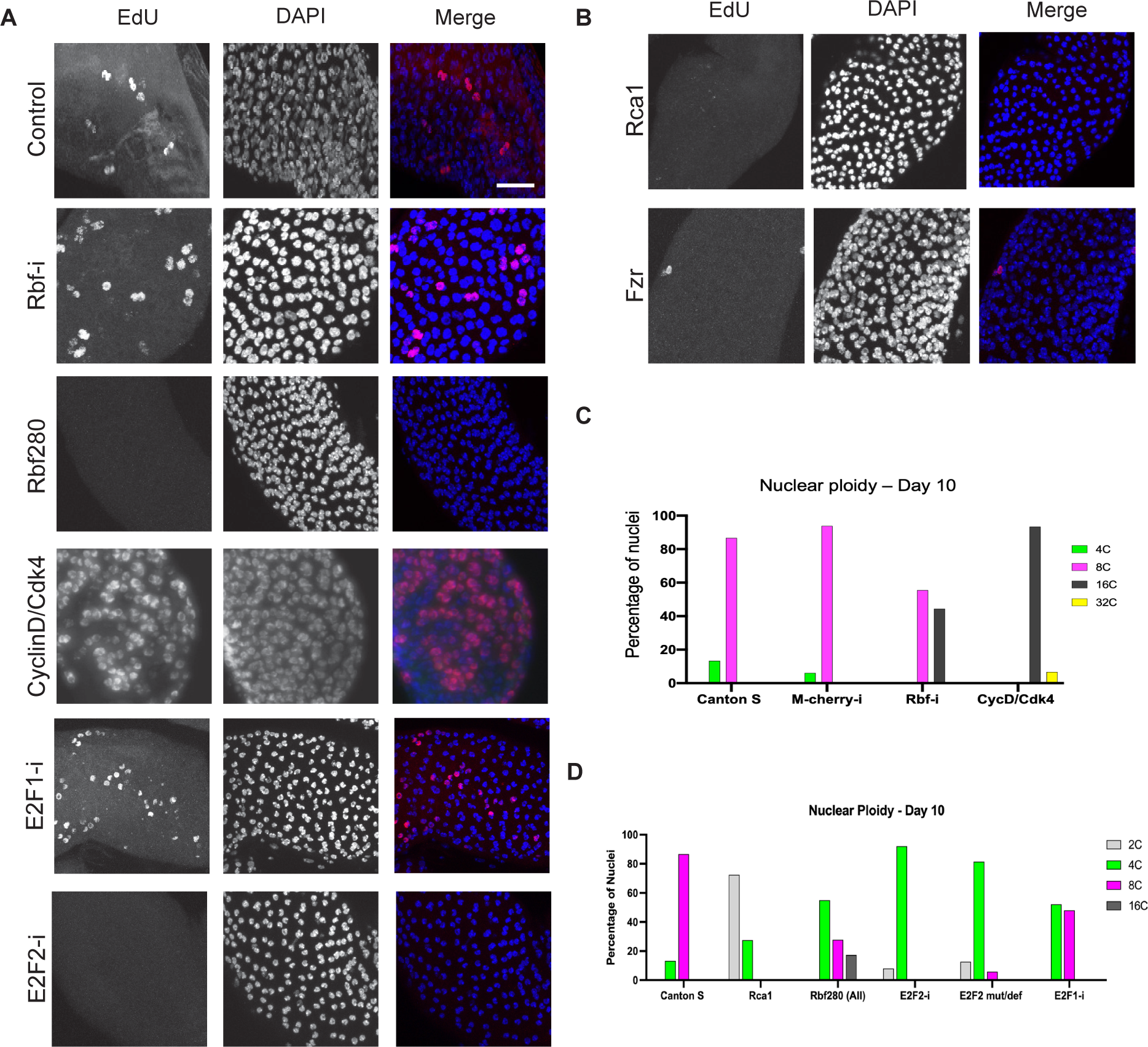
The E2F/Rbf network and APC/C activity oscillations regulate adult stage main cell endocycles. A: EdU incorporation on DOE while manipulating the E2F/Rbf network. Control is Prd-Gal4 males without genetic manipulation. Genotypes used to disrupt the E2F/Rbf network were crossed to Prd-Gal4 and are indicated in the figure. B: EdU incorporation on DOE while manipulating APC/C oscillations. Genotypes used to disrupt the APC/C oscillations were crossed to Prd-Gal4 and are indicated in the figure. C: Quantification of nuclear DNA content for Day 10 virgins represented as percentage of nuclei measured. This graph shows manipulations that increased ploidy. D: Quantification of nuclear DNA content for Day 10 virgins represented as percentage of nuclei measured. This graph shows manipulations that decreased ploidy. Scalebar: 20 microns

We next examined the role of E2Fs, E2F1 and E2F2, in AG main cell endocycling. When E2F1 is knocked down in the AG, we observe widespread EdU incorporation (Figure 3A). Importantly, this result is coupled with nuclear DNA content measurements that are, slightly less than, but similar to wildtype. This suggests that the observed increase in EdU labeling when E2F1 is inhibited likely occurs by slowing or prolonging the endocycle rather than the presence of extra endocycles per cell. Consistent with this, at 10 days we see several nuclei with an intermediate DNA content between 4C and 8C (Figure 3D) suggesting that some nuclei with reduced E2F1 may fail to complete the adult post-eclosion endocycle for several days. Strikingly, when E2F2 was depleted via RNAi we find EdU incorporation on the day of eclosion was nearly abolished in all animals analyzed (Figure 3B). Additionally, DNA content of main cell nuclei was measured on Day 10 and found to be only 4C (Figure 3D), suggesting that reducing E2F2 prevents endocycling and that E2F2 may be necessary for the post-eclosion endocycle. To confirm this, we performed DNA content measurements on main cells of E2F2 null mutant/deficiency males at day 10, which also show a loss of the 8C population (Figure 3D). This demonstrated that loss of E2F2 prevents the post-eclosion endocycle, possibly in a manner similar to that reported in other endocycling tissues (Maqbool et al., 2010; Weng et al., 2003). Altogether our data shows that the E2F/Rbf network is a critical regulator of proper endocycling in main cells and reveals an essential role for E2F2 in either directly or indirectly promoting the first adult endocycle in this tissue.

In addition to oscillations of E2F activity, oscillations of the Anaphase Promoting Complex/Cyclosome (APC/C) are also required for proper endocycle progression in other tissues like the ovary and salivary gland (Narbonne-Reveau et al., 2008). The APC/C is a ubiquitin ligase that targets many cell cycle proteins for destruction and in the endocycle APC/C activity oscillations are essential for degradation of the replication inhibitor Geminin (Zielke et al., 2008). To determine whether inhibition of APC/C oscillations would also disrupt proper endocycling in the main cells, we overexpressed Regulator of Cyclin A1 (Rca1; in mammals Emil1), an inhibitor of APC/C activity, in the AG. When we inhibit the APC/C in this manner, EdU labeling on the day of eclosion in suppressed (Figure 3B) and main cells have a nuclear DNA content of 2C on day 10, resulting in bi-nucleated main cells with a DNA content as low as 4C (Figure 3D). This suggests that blocking the APC/C effectively prevents the pupal endocycle as well as subsequent endocycles in the adult tissue. We observe a similar loss of EdU incorporation on the DOE with the knockdown of the APC/C regulatory subunit *Fizzy-related* (*Fzr*, ortholog of FZR1, also known as Cdh1), demonstrating APC/C^Fzr^ activity is required (Figure 3B). This establishes that E2F and APC/C activity oscillations are critical for proper endocycle progression in the *Drosophila* AG main cells and that the AG maintains plasticity regarding the timing and progression of the post-eclosion endocycle and final cellular DNA content.

### Juvenile hormone signaling is required for adult accessory gland endocycling and fertility

Juvenile Hormone (JH) signaling has been shown to play an important role in regulating adult reproduction for many types of insects, including *Drosophila*. Previous work in migratory locusts show this hormonal signaling can act upstream of the cell cycle to promote polyploidy in the fat body that is critical for proper vitellogenesis (Wu et al., 2016, 2018, 2020a) and JH signaling promotes AG growth and protein synthesis specifically in *Drosophila* (Herndon et al., 1997; Shemshedini et al., 1990; R. Yamamoto et al., 2013). There is a pulse of JH that begins just prior to eclosion, which peaks shortly after eclosion (Bownes and Rembold, 1987). We therefore hypothesized that JH signaling may play a role in regulating endocycling on the day of eclosion in AG main cells. To examine this, we disrupted JH signaling in the AG by using *prd*-Gal4 to express RNAi for JH nuclear receptors, *germ cell-expressed bHLH-PAS* (*Gce*), *Methoprene tolerant* (*Met*), and their co-activator *Taiman* (*Tai*). *Gce*, *met*, and *tai* have all been shown to be expressed in the adult AG (A. Baumann et al., 2010; A. A. Baumann et al., 2017; Jindra et al., 2015). RNAi to JH receptor *Gce* and co-activator *Tai* each abolish EdU incorporation in the AG on the day of eclosion in 1 hour ex vivo experiments (Figure 4A). For both genetic manipulations, main cells also have reduced ploidy in day 10 virgins with a total loss of the 8C population that is observed in controls (Figure 4B), suggesting that JH signaling through *Gce* and *Tai* is necessary for proper main cell endocycling.

**Figure 4:**
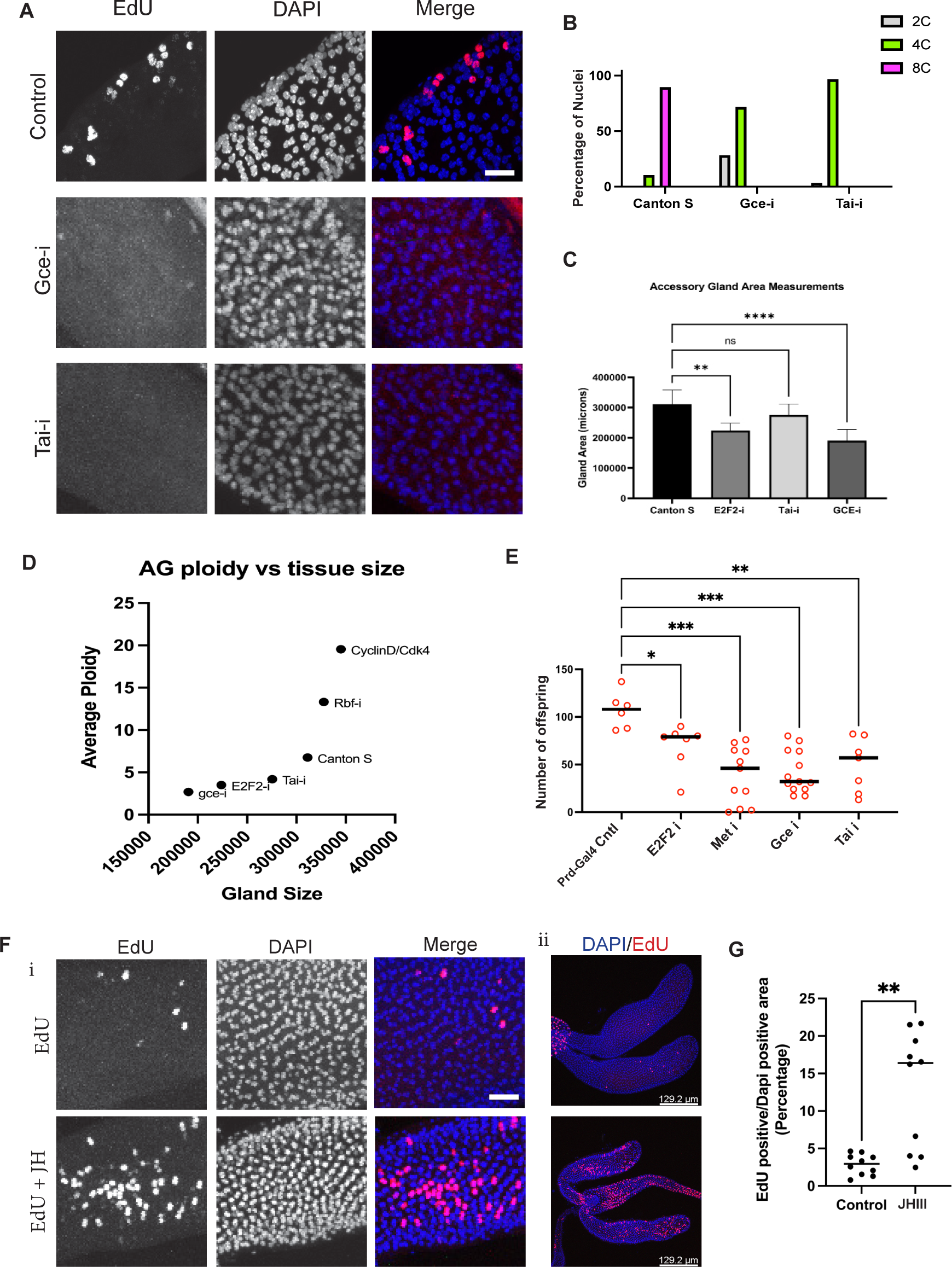
Juvenile hormone signaling is required for adult accessory gland endocycling and fertility. A: EdU incorporation assays on DOE while manipulating juvenile hormone signaling. The control is Prd-Gal4 males without genetic manipulation. Genotypes used to disrupt juvenile hormone signaling were crossed to Prd-Gal4 and are indicated in the figure. B: Quantification of nuclear DNA content for Day 10 virgins represented as percentage of nuclei measured. Disruption of juvenile hormone signaling shows decreased main cell ploidy. C: Accessory gland area measurements for 10 day adult accessory glands for indicated genotypes. Disruption of main cell endocycling shows decreased gland area. D: AG nuclear ploidy vs AG gland size. Accessory gland size scales with main cell ploidy. E: Fertility assay showing the total number of offspring that result from males of indicated genetic manipulations crossed to wild-type Canton S virgin females. Males that contain accessory glands with reduced nuclear ploidy show reduced number of offspring. F: EdU incorporation on DOE with and without the addition of juvenile hormone III. Labeling was done on 1 hour post eclosion animals, for 1 hour, ex vivo. (i) shows 63x mag (ii) shows 20x mag of the same tissues. G: Quantification of EdU incorporation from (F). Data shown is DAPI positive area that is also EDU positive plotted as a percentage. In a subset of tissues examined, the addition of JH-III causes increased levels of EdU incorporation. Scale: A: 15 microns F i: 15 microns F ii: 129 microns Statistical Analysis: C: Ordinary one-way ANOVA, ** 0.0013, **** <0.0001 E: Ordinary one-way ANOVA, *., **., ***, G: Welch’s T-test

Conservation of main cell endocycling on the day of eclosion across *Drosophila* species suggests there may be an important biological role for this phenomenon. We next asked how blocking the endocycle on the day of eclosion affects this tissue. We report a decrease in the overall size of the adult gland when JH signaling is inhibited (Figure 4C), suggesting that early gland growth is driven, in part, by endocycles. Importantly, we also see a reduction of tissue size in *E2F2* RNAi conditions (Figure 4C) which suggests the reduced gland size phenotype is caused by the loss of the endocycle and is not specific to the disruption of JH signaling. Additionally, we observe that gland size appears to scale with ploidy when manipulations that decrease DNA content take place (Figure 4D), where the manipulations that disrupt ploidy to the greatest extent are smaller than tissues with manipulations that disrupt ploidy to a lesser extent. We see that males with reduced cellular DNA content and reduced overall tissue size show a statistically significant reduction in fertility (Figure 4E). This suggests that the tissue-wide endocycle that occurs shortly post-eclosion is critical to establish optimal cellular ploidies and tissue size, which may drive proper gland development and function.

JH signaling is necessary to drive the post-eclosion endocycle in the adult AG (figure ***). Therefore, we next asked if JH is sufficient to induce endocycle entry. To test this hypothesis, we collected animals within an hour of eclosion and performed an ex vivo Ringers+EdU labeling protocol where we added ectopic synthetic Juvenile Hormone (JH III), or vehicle only (acetone). We observed an increased level of EdU incorporation in the adult accessory gland with JH III (Figure 4F,G) compared to controls, suggesting that JH is sufficient to push the AG main cells of newly eclosed males to enter S-phase.

### The *Drosophila* accessory gland undergoes progressive cell cycle remodeling throughout development to create a uniquely structured polyploid tissue

The development of the AG involves cell cycle remodeling from canonical proliferation into variant cell cycles, coordinated with cellular terminal differentiation. The cell cycle remodeling results in progressive truncations of the cell cycle, first by limiting cytokinesis to result in bi-nucleation, followed by endocycles which lack M-phase altogether. Here we have shown that the final post-eclosion endocycle exhibits a further truncation of S-phase resulting in under-replicated heterochromatin. This process of progressive cell cycle truncation led us to wonder whether additional cell cycle truncations may take place during gland development.

Ring canals are created when cytokinesis is truncated and the cytokinetic furrow does not fully close and is instead stabilized. The resulting actin-rich structure creates an opening between cells through which cytoplasm is shared (McLean & Cooley, 2014). It has been suggested that ring canals may help to balance protein levels in cells that have uneven transcription by allowing them to share intracellular molecules, which may be beneficial for this highly secretory tissue. Using a line containing ubiquitously expressed Pavarotti (Pav) tagged with GFP (Pav-GFP), we and others (Eikenes et al. 2013) observe localization on the cell membrane of main cells in the adult AG (Figure 5A). Pavarotti is a kinesin like protein that has been previously shown to be stably localized to ring canals in other *Drosophila* tissues, most notably the follicle cells (Airoldi et al., 2011).

**Figure 5:**
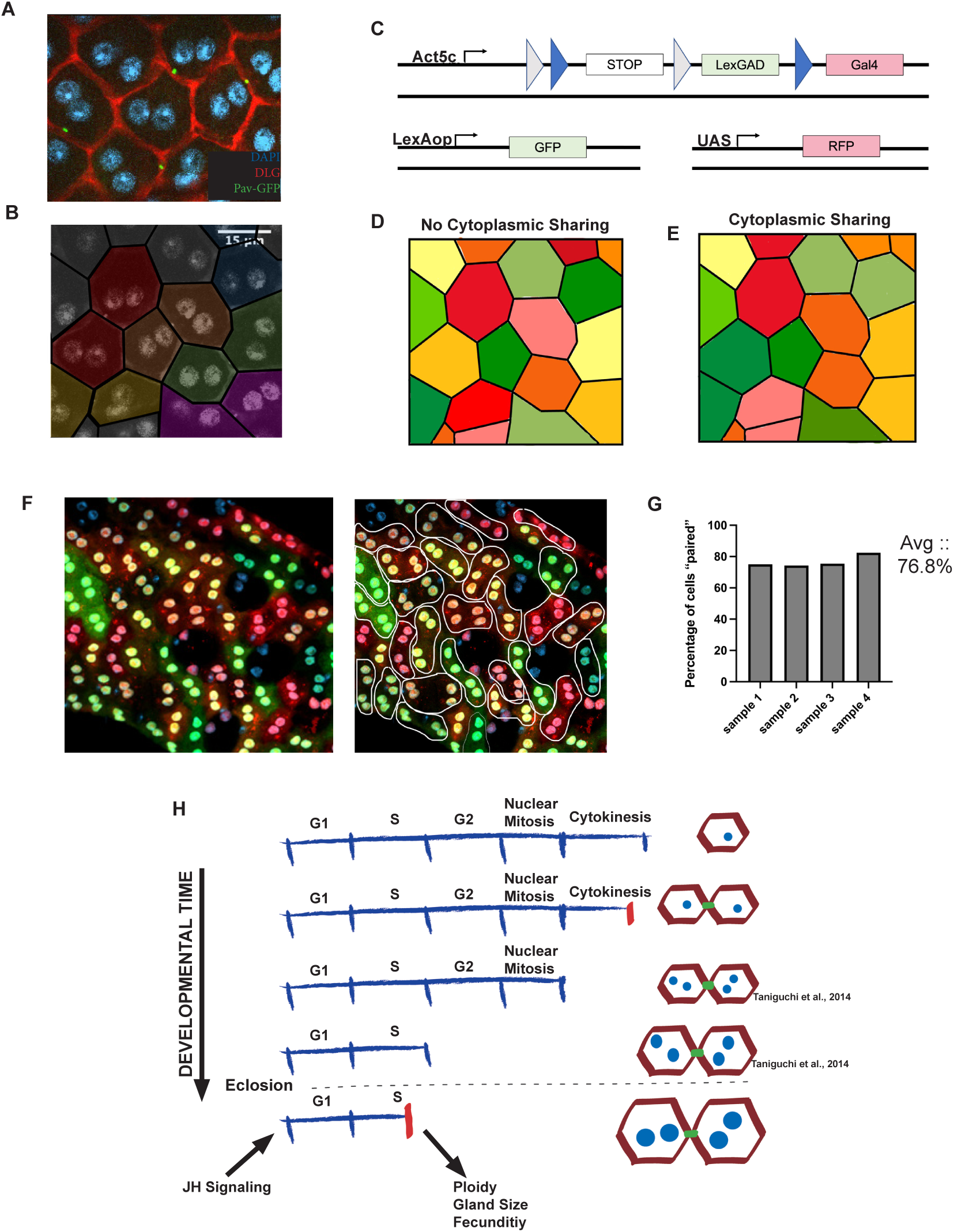
The *Drosophila* accessory gland undergoes progressive cell cycle remodeling throughout development to create a uniquely structure polyploid tissue. A: Pavarotti-GFP localization in adult tissues. Each cell contains 1 foci of Pav-GFP located medially on a shared membrane space. B: Psuedocoloring of cells sharing a Pav-GFP foci from a microscopic image of the adult accessory gland. Psuedocoloring creates a pattern of pairs of main cells. C: Schematic of experiment using CoinFLP (modified from Bosch et al., 2015). Flies also contain a heatshock induced flippase enzyme to allow for temporal control of CoinFLP activation. D: Cartoon diagram of what CoinFLP labeling would look like in an adult AG when flipped after gland development if no cytoplasmic sharing is present. E: Cartoon diagram of what CoinFLP labeling would look like in an adult AG when flipped after gland development if cytoplasmic sharing is present. F: Representative image of CoinFLP in the adult AG. Flipping occurred in the adult stage after gland development has occurred. Animals were dissected 1 day post induction so that fluorescence levels were given time to equilibriate if cytoplasmic sharing is present. G: Same image as F – with outline showing “cells that are paired” in fluorescence levels. G: Quantifications of number of cells “paired” in fluorescence levels in all samples analyzed. The average is 76.8% of cells, which is higher than expected for randomized flipping without cytoplasmic sharing. H: Model of cell cycle variants occurring throughout AG development. Red lines on the linearized cell cycle represent a truncation of that cell cycle phase. Green structures on the cellular models represent ring canal structures. This study has added the upstream, final mitotic cell cycle that creates ring canals/cytoplasmic sharing and the post-eclosion truncated endocycle that is dependent on JH signaling and required for proper gland development and fertility.

Interestingly, the location and number of Pav-GFP foci in the adult AG displayed a distinct and reproducible pattern. Only one Pav-GFP focus is present on the membrane of each main cell and is located centrally at bicellular junctions. Two neighboring main cells can have a single ring canal between them, but they will have no other ring canals between themselves and any other cells of the AG. This pattern suggests that Pav-GFP delineates ring canals that arose from a truncated, penultimate truncated cell cycle, which generates sister cells prior to the binucleation event during AG development (Figure 5B, psuedocoloring of sister cells based off true Pav-GFP localization). Importantly, Pav-GFP is not seen on the membrane of secondary cells, suggesting that these cells may not communicate via shared cytoplasm with their neighboring main cells.

To test whether the pattern of Pav-GFP was indicative of functional ring canals allowing cytoplasm exchange, we used a genetic tool that allows multi-color labeling of polyploid cells. CoinFLP, is a genetic tool in which a single chromosome can flip out a cassette to express either LexGAD or Gal4, inducing either green or red fluorescent protein when flippase enzyme is present (Figure 5C) (Bosch et al., 2015). By using a heatshock-induced flippase we limited expression of flippase to adult tissue. Due to the large number of chromosome copies in each 8C nucleus and 16C cell, we observe multiple combinations of CoinFLP flipping, creating cells with varying levels of green, yellow, orange, and red flourescence. If no cytoplasmic sharing occurs, the range of colors will vary cell by cell regardless of time passed since flipping (Figure 5D). However, if cytoplasmic sharing does occur between sister-cells we would see the color between the two cells equalize over time, creating a pattern of pairs of main cells with similar colors (Figure 5E). When we induce CoinFLP in the adult tissue and allow time for cytoplasmic sharing to occur, we see what appears to be paired main cells, immediately next to each other of the same color indicating cytoplasmic sharing in the adult gland (Figure 5F). When we quantify the number of cells per frame that appear to be color sharing across samples, the average is 76.8% (Figure 5G). Together, these data suggest that main cells form ring canals from a truncated cytokinesis in the penultimate mitotic cell cycle. Our work, combined with the work of others, supports a model for developmentally regulated variant cell cycles in *Drosophila* AG main cells that involves a progressive truncation of the canonical mitotic cell cycle to create a uniquely polyploid tissue (Figure 5H).

## Discussion

Here we present data supporting a model for progressive cell cycle truncations in the main cells of the *Drosophila melanogaster* accessory gland. Our data suggest a penultimate mitotic cell cycle occurs in the pupa accessory gland, prior to binucleation, with a partially truncated cytokinesis that forms ring canals in paired sister main cells. Around 50-60 hours APF, all main cells undergo a further truncated cycle in which nuclear mitosis proceeds but cytokinesis does not occur, leading to bi-nucleation (Taniguchi et al. 2014). Then at 70-80h APF, the final cell cycle during metamorphosis is an endocycle completely lacking mitosis (Taniguchi et al. 2014). Here we also provide evidence of an additional endocycle in the adult gland that under-replicates heterochromatic regions of the main cell nuclei via a truncated S-phase within the first 5 hours of eclosion. We propose this additional endocycle is important for reproductive maturity to increase gland size and protein synthesis to support male fertility.

### Endocycling in adult accessory gland maturation and function

A wave of DNA replication occurs in most, if not all, main cells of the accessory gland shortly after eclosion. Using approaches to reduce or block the post-eclosion endocycle, we show this endocycle is important for proper gland growth, maturation, and male fecundity. Similar to other endocycling tissues, APC/C activity oscillations and the E2F/Rb network is critical for the endocycling status of the adult AG main cells. While the proper timing of the endocycle in main cells requires E2F1 activity oscillations, the endocycle can be nearly completely blocked by inhibiting the APC/C or loss of E2F2 function, suggesting this tissue is particularly sensitive to the threshold levels of these endocycle regulators. Tissue-specific knockdown of E2F2 by RNAi phenocopies the loss of gland endocycling in an *e2f2* null mutant background, including reduced fertility, suggesting defects in somatic male reproductive tissues may also contribute to male fertility defects in cell cycle mutants.

Juvenile Hormone has previously been shown to promote AG growth (Herndon et al., 1997; Shemshedini et al., 1990; T. G. Wilson et al., 2003; K. Yamamoto et al., 1988). Here we show that endoreplication, gland size, and overall fertility are affected when this signaling pathway is disrupted. This opens a line of inquiry on what downstream cell cycle components juvenile hormone directly or indirectly regulates in the *Drosophila* AG. Prior work in migratory locusts has demonstrated that Juvenile Hormone signaling can impact levels of key cell cycle regulators in endocycling fat body cells to support vitellogenesis. Juvenile Hormone signaling can impact expression of the E2F1 transcription factor itself, the replication licensing component cdc6 and essential DNA replication factors such as MCMs and Orcs through direct transcriptional regulation via the JH receptor complex or indirect signaling pathways (Guo et al., 2014; Wu et al., 2018, 2020b). Whether the impacts of JH signaling on the endocycle in the *Drosophila* AG are through direct transcriptional regulation of E2F factors and DNA replication or more indirect signaling will need to be investigated. The AG provides a novel cellular context to examine how JH signaling impacts the endocycle with functional tissue assays of growth and fertility. This will also allow for further investigations to decipher whether JH signaling promotes biosynthesis indirectly through the endocycle, or also more directly through cellular growth pathways (Øvrebø et al., 2022).

Another hormonal signaling pathway, the ecdysone steroid signaling pathway has previously been shown to impact endocycling in the *Drosophila* accessory gland secondary cells (Leiblich et al., 2019; Sekar et al., 2023) although additional work also suggests it affects AG cell survival, growth and male fertility in the gland more broadly, including in main cells (Sharma et al., 2017). In secondary cells, the Ecdysone receptor can act independent of the steroid hormone to promote the cell cycle through the RB/E2F pathway, although due to feedback loops in cell cycle regulation with Cyclins, the RB/E2F pathway acts both upstream and downstream of the ecdysone receptor in regulating the cell cycle of secondary cells. Our work adds an additional hormonal signaling axis to the picture of AG maturation, one that regulates cycling in the predominant AG cell type of main cells, affects gland size, and is essential for proper male fertility. Additional work will be required to investigate the interactions between the ecdysone and juvenile hormone signaling pathways, which have previously been shown to have both a cooperative and an antagonistic relationship depending upon cellular context (Gao et al., 2022; Liu et al., 2018; Zipper et al., 2020).

### The Accessory Gland as a model to study multiple, successive cell cycle truncations

During accessory gland development, the canonical mitotic cell cycle is remodeled via cell cycle truncations into variant cell cycles throughout late metamorphosis and early adulthood. These variant cell cycles are precisely timed and occur relatively synchronously throughout the tissue suggesting a tissue-wide level of developmental cell cycle control. This strict regulation of timing brings forth a unique opportunity in which we can begin to manipulate these truncations one at a time, or even in a pair-wise manner, to see the resulting effects on the tissue structure and function. Previous work has shown that the loss of binucleation can affect the mechanical abilities of this tissue which alter reproductive success (Taniguchi et al., 2018). Here we examine previously unappreciated cell cycle plasticity in the adult tissue to understand how cell cycle variants after the mitotic to endocycle switch to impact gland growth and function (Figure 5H -model). Future work will be aimed at deciphering the cell signaling pathways that remodel the cell cycle in this tissue to understand the advantages of a polyploid and binucleate state. Binucleate cells are found in the mammalian urothelium, mammalian cardiac muscle and secretory cells of the mammary gland, where they are also often polyploid (Rios et al., 2016; Swift et al., 2023; Wang et al., 2018). Of note these tissues are flexible, either contracting repeatedly or responding to lumen filling, and exhibit unusual elasticity. We hope to use the *Drosophila* accessory gland to understand how the polyploid and binucleate state impacts epithelial cell biology.

## Acknowledgements

We thank members of the Buttitta and Taichman Labs for advice and discussions, especially Ajai Pulianmackal, Dr. Frank Cackowski and Dr. Kenji Yumoto. We thank Pusparanee Hakim and Ce Wang for help with early experiments characterizing the fly AG and Yukiko Yamashita for helpful discussion. This work was supported by a Prostate Cancer Foundation Challenge Award (16CHAL05) and an American Cancer Society Scholar Award (RSG-15-161-01-DDC). A. Box was supported by the University of Michigan Organogenesis Predoctoral Training Grant (NIH T32 HD007505) and the Eunice Kennedy Shriver National Institute of Child Health and Human Development of the National Institutes of Health (NIH F31 HD103430-01). S.J. Church was supported by a University of Michigan Rackham Merit Fellowship and S. Nandakumar was supported by a University of Michigan Barbour Scholar Award. Work in the Buttitta Lab is also supported by the National Institutes of Health (NIH R01 GM127367) and work in the Taichman Lab is supported in part by the NCI (NIH P01 CA093900).

## Materials and Methods

### Fly stocks

Canton S

*D. simulans* (Gift from Patricia Wittkopp Lab)

*D. yakuba* (Gift from Patricia Wittkopp Lab)

*D. pseudoobscura* (Gift from Patricia Wittkopp Lab)

*D. willistoni* (Gift from Patricia Wittkopp Lab)

*D. virilis* (Gift from Patricia Wittkopp Lab)

Prd-Gal4 (BL#1947): w[*]; Prd-Gal4/TM3, Sb[1] w; +; Prd-Gal4, UAS-His2AV::RFP

Rbf-i: W; if/cyo (lacz); UAS-Rbf RNAi

Rbf280 (BL#50748): w[*]; P{w[+mC]=UAS-Rbf.280}3/TM3, Sb[1]

Dk4: y,w,hs-flp; UAS CycD, UAS-Cdk4/CyO-GFP

E2f1-i: y,v; UAS-E2F1 RNAi

E2f2-i (BL#36674): y[1] sc[*] v[1] sev[21]; P{y[+t7.7] v[+t1.8]=TRiP.HMS01562}attP2

Rca1: y,w, hs-flp; Pin/CyO-GFP; UAS-HA-Rca1/TM6B

Fzr-i: y,w,hs-vlp; UAS-Rap^IR^/ CyO-GFP; UAS-E2F1, UAS-Dp/ TM6B

Ek2: y,w, hs-flp; +; UAS-CycE,UAS-Cdk2/TM6B

Gce-i (BL#26323): y[1] v[1]; P{y[+t7.7] v[+t1.8]=TRiP.JF02097}attP2

Tai-i (BL#32885): y[1] sc[*] v[1] sev[21]; P{y[+t7.7] v[+t1.8]=TRiP.HMS00673}attP2 Met-i (BL#26205): y[1] v[1]; P{y[+t7.7] v[+t1.8]=TRiP.JF02103}attP2

Mcherry-i (BL#35785): [1] sc[*] v[1] sev[21]; P{y[+t7.7] v[+t1.8]=VALIUM20-mCherry}attP2

Pav-gfp: (Gift from Yukiko Yamashita Lab)

CoinFLP: P{CoinFLP-LexA::GAD.GAL4}

CoinFLP reporter line: y,w,hs-flp; lex-op-nls-gfp/cyo; uas-nls-rfp/tm6b

### Fly rearing and mating

All flies were raised and kept at room temperature (23°C) on Bloomington Cornmeal food unless otherwise noted. All experiments used virgin males unless otherwise noted: males were collected on the day of eclosion (1-3 hours post eclosion) and aged for indicated times in vials containing no more than 7-10 males per vial. For experiments with mated animals: males and females were collected as virgins, on the day of eclosion, and were kept at an approximate 1:1.5 ratio for indicated times unless otherwise stated.

### Tissue fixation and staining

Accessory glands were dissected in 1X PBS and were fixed in 4% PFA+1xPBS for 30 min at room temperature while rocking. Tissues were rinsed with 1xPBS+0.1%Triton-X twice for 10 minutes. Tissues were further permeabilized in 1xPBS+1.0%Triton-X for 30 minutes at room temperature while rocking. Tissues were rinsed with PAT for 10 min and primary antibodies diluted in fresh PAT were incubated at room temperature rocking overnight. Tissues were rinsed twice for 10 minutes in 1xPBS+0.1%Triton-X. Tissues were pre-blocked in PBT-X+2%NGS for 10 minutes. Secondary antibody was added to fresh PBT-X+2%NGS and tissues were incubated overnight rotating at room temperature. Tissues were rinsed twice in 1xPBS+0.1%Triton-X before incubating in DAPI (1µg/ml) for 10 min. Tissues were rinsed thoroughly with 1xPBS+0.1%Triton-X before being mounted with Vectashield. Mounting was done using “wells” created by 1 layer of clear nail polish so that tissue would not be flattened and luminal space remained intact.

The following antibodies were used in this study: Mouse anti DLG (DHSB) 1:500, Rabbit anti PH3 (Millipore #06-570) 1:1000, mouse anti PH3 (Cell Signaling #9706) 1:1000.

### Timing of EdU uptake

In this study, we use blue food coloring in our sucrose solution to ensure that flies we are analyzing have taken up the solution. To time when flies are eating post-eclosion we looked at the flies every 30 minutes to check for “blue bellies” and graphed this data in Prism Graphpad to report the percentage of animals that do *<u>not</u>* exhibit blue bellies and therefore have not eaten at each time point.

### EdU labeling

Click-IT Plus EdU AlexaFluor-555/488 Imaging kits were used as directed (Life Technologies).

For labeling on the day of eclosion, accessory glands were dissected between 1-3 hours post-eclosion and immediately placed into Ringers solution containing 0.1 mM EdU for 1 hour prior to fixation.

For long term adult labeling, animals were fed 1 mM EdU in 10% sucrose with blue food coloring for the indicated amounts of time. EdU/sucrose mixture was placed on Whatman paper in empty vials and was changed every 2-3 days to control for contamination. We also performed feeding with 1 mM EdU in Cornmeal food with blue food coloring for up to 6 days and obtained similar results. The blue food coloring allows for visualization of which animals have ingested the sucrose solution.

For juvenile hormone incubation: Juvenile Hormone III (Sigma Aldrich) was added to a Ringers/0.1mM EdU solution for ex-vivo labeling. JH-III was reconstituted in acetone as a 50x master mix and used at .1ug/.5ul concentration. The control experiment contained the same amount of acetone as the JH induction, without the addition of the hormone. Males were collected from 0-1 hours post eclosion and were labeled with EdU for 1 hour before fixation.

### Measurements/DAPI quantifications

Fluorescent images were obtained using a Leica SP5 confocal, Leica SP8 confocal or Leica DMI6000B epifluorescence system.

Gland measurements were completed by using brightfield microscopy on the Leica DMI6000B epifluorescence system. Max projections were exported into Image J, where measurements were obtained, unless otherwise specified measurements are reported in microns. Due to the measurements being on projection, these measurements are not of gland volume, but of “2D area.”

Discs large (DLG) antibody staining was used for all cell size measurements. Due to the apical localization of DLG, cell size measurements reported here are not of the volume of a cell, but rather a measurement of the apical area. Measurements reported here are taken mid-lobe and are only of main cells to ensure cell type differences are not confounding the measurements. Image J was used to obtain measurements of cells in microns and measurements were transferred to Prism for statistical analysis.

Nuclear area measurements were done similarly to cell size (described above) using DAPI signal to delineate nuclei. The measurements reported here are taken mid-lobe and include main cells only.

To obtain ploidy measurements, we modified a protocol from the Losick Lab (Grendler et al., 2019) for this tissue. In brief: DAPI was carefully imaged using Leica SP5 confocal system and intensity quantifications were measured using ImageJ. Raw Integrated Intensity measurements were used for the binucleated cells from the mid-lobe region of the accessory gland and the mononucleated, basally located, diploid muscle nuclei of the ejaculatory duct where Paired-gal4 is not expressed. The average background fluorescence was subtracted from each intensity measurement to get the corrected intensity. Intensity for haploid DNA content was deduced by dividing the average intensity of the basally located, ejaculatory duct muscle nuclei in half. Ploidy of the accessory gland nuclei was established using the following binning: 2N (1.9-2.9), 4N (3.0-6.9), 8N (7-12.9), 16N (13.0-24.9), 32N (>24.9).

### Flow Cytometry

Flow cytometry was done on a nuclear prep of the AG modified from previous studies (Majane et al., 2022; McLaughlin et al., 2022). Briefly, tissues were dissected in cold S2 media and kept on ice. S2 media was removed, lysis buffer was added and tissues were transferred to cold dounce homogenizers. Tissues were dissociated using 20 strokes with the loose and 40 strokes with the tight pestle. Samples were ran through 100 and 40 micron filters and a small amount of S2 was added to quench the lysis reaction. Samples were spun on a tabletop nutator to create a nuclear pellet. S2/lysis media was removed from pellet of nuclei and pellet was resuspended in fresh S2 media with Vybrant DyeCycle Violet stain (Invitrogen) at 1:100 and left to incubate on ice for 10 minutes. Samples were spun again and S2/DyeCycle Violet media was replaced with fresh S2. Samples were lightly vortexed just prior to running on the Attune Flow Cytometer.

To plot the peak shift between control animals and cyclinE/cdk2 overexpression animals: Due to the small nature of the shift we see on the histograms, we wanted to ensure that the shift was not due to a difference in sample prep between our control and our genetic manipulation samples. Thus, we performed the experiment using two genotypes in parallel. Each experimental setup used a fly line in which only the accessory gland nuclei were labeled with RFP using a Prd-Gal4, UAS-His2AV:RFP. This line was crossed to either w^1118^ or w; UAS-CyclinE, UAS-cdk2. For each cross, we prepared the accessory glands and ejaculatory ducts in the same vial, so that we could sort the positively labeled accessory gland nuclei from the RFP negative ejaculatory duct cells. We aligned the non-genetically manipulated ejaculatory duct histograms from each cross to ensure that sample prep was equal amongst the control and the genetically manipulated sample. We show the AG plots only.

### Fertility

We modified a previously established method (Herrera et al., 2021). In brief: virgin males of stated genotypes and Canton S virgin females were collected between 16-24 hours prior to starting assay. 2 Canton S virgin females and 1 virgin male of stated genotype were mated for two days at 25°C. Total offspring arising from the individual crosses were counted. Counts were done on pupa and emerging adults to assure that there were no viability issues within the pupal stage.

### CoinFLP

CoinFLP experiments in the accessory gland were performed in the adult only. The animals were heat-shocked for 20 minutes in a 37C water bath on day 9 post eclosion. AGs were dissected 24 hours post heat shock, fixed, counterstained with DAPI and fluorescence levels were observed via confocal microscopy (Nandakumar & Buttitta, 2023).

### Statistical Analysis

All statistics were run with Prism Graphpad. Analysis performed for each experiment is reported in the figure legends.

---

**Supplemental Figure 1:**
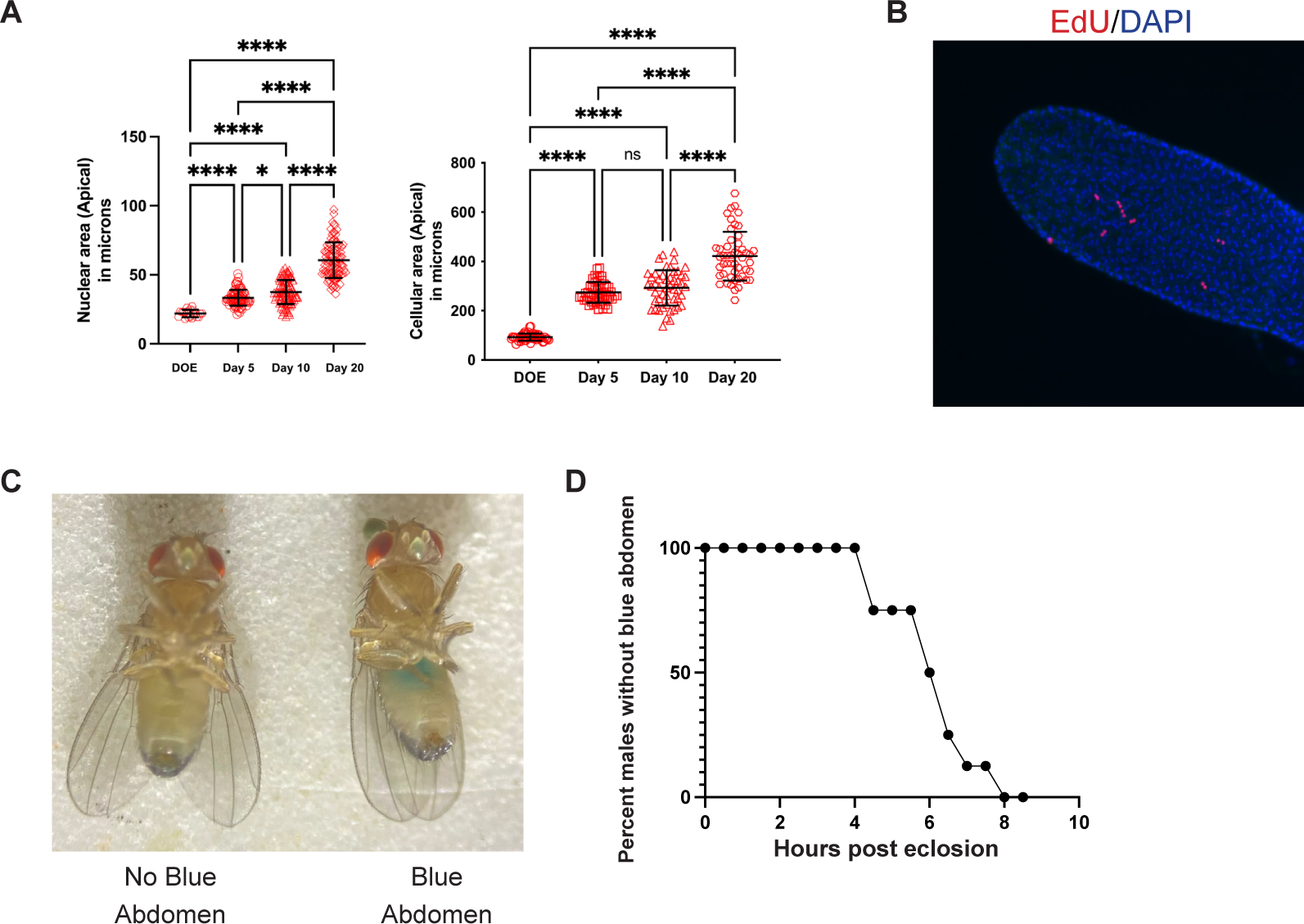
Endocycling in the adult accessory gland. A: Nuclear and cellular area measurements with age in virgin adult *D. melanogaster* accessory gland B: EdU incorporation in a Day 10 virgin male. EdU+Sucrose was fed to the animals from ∼6 hours post eclosion until Day 10. C: Visualization of blue food coloring through the abdomen. Blue food coloring is added to all feeding assays to ensure that the animals analyzed have ingested the solution. The photo shown here is of two flies from the same feeding vial around 4 hours post-eclosion – showing that some animals have taken in the solution at this point, while some have not. D: Quantification of the timing that flies take in solution via feeding. Graph is showing the percentage of flies without blue bellies over time. Statistical Analysis: A: Ordinary one way ANOVA *0.0226, **** <0.0001

**Supplemental Figure 2:**
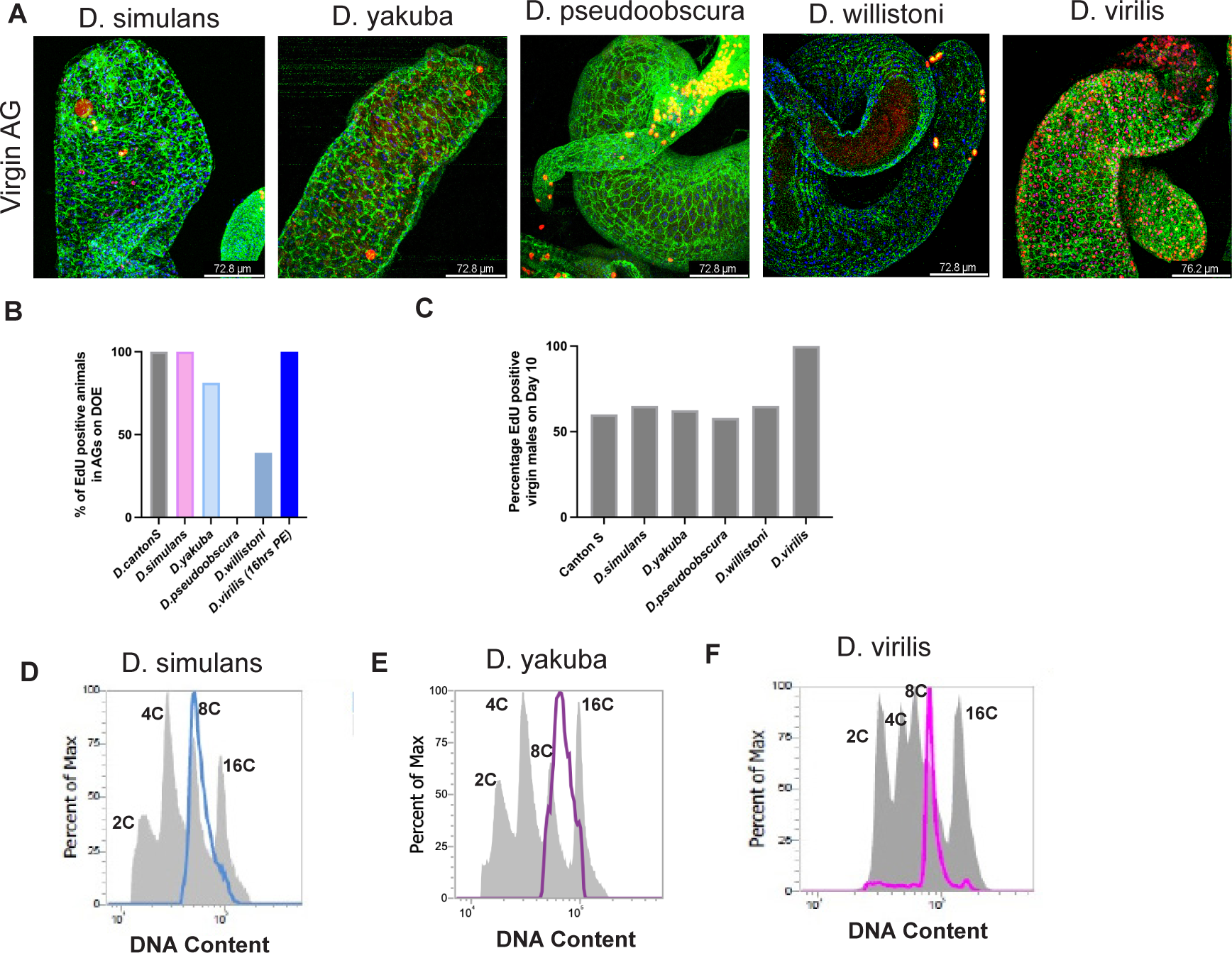
Endocycling and polyploidy of main cells is conserved in other *Drosophila* species. A: EdU incorporation in Day 10 virgin males for multiple *Drosophila* species. EdU+Sucrose was fed to the animals from ∼6 hours post eclosion until Day 10. EdU incorporation is conserved. B: Quantification of EdU labeling on DOE in multiple *Drosophila* species. Data shown as percentage of animals that are positively labeled with EdU during a 1 hour ex vivo labeling on DOE. C: Quantification of EdU labeling on Day 10 in multiple *Drosophila* species. Data shown as percentage of animals that are positively labeled with EdU over a 10 day feeding. D, E, F: Flow cytometry histograms of nuclear DNA content of multiple *Drosophila* species.

**Supplemental Figure 3:**
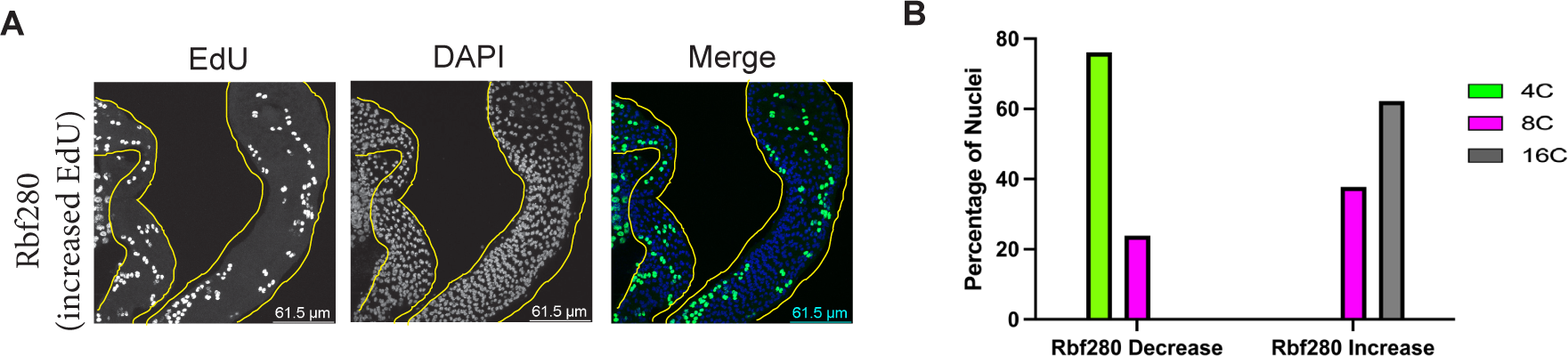
Further characterization of endocycling in the adult accessory gland. A: EdU incorporation on DOE. A subset of animals expressing Rbf280 show increased EdU incorporation as shown here. AGs are outlined in yellow. B: Quantification of nuclear ploidies at Day 10 post eclosion represented by percentage of nuclei. Here, we separate the two phenotypes from RBF280 expression. This data is shown combine in Figure 3.

